# Multilevel selection in groups of groups

**DOI:** 10.1101/624718

**Authors:** Jonathan N. Pruitt, David N. Fisher, Raul Costa-Pereira, Noa Pinter-Wollman

**Author notes:** **Denotes corresponding author**.

## Abstract

Natural selection occurs at many levels. We evaluated selection acting on collectives at a level of multilevel selection analysis not yet quantified: within and between clusters of groups. We did so by monitoring the performance of natural colonies of social spiders with contrasting foraging aggressiveness in clusters of various sizes. Within-clusters, growth rates were suppressed when colonies were surrounded by more rival groups, conveying that competition is greater. When colonies were surrounded by few rivals, the more aggressive colonies in a cluster were more successful. In contrast, relatively non-aggressive colonies performed better when surrounded by many rivals. Patterns of selection between-clusters depended on the performance metric considered, but cluster-wide aggressiveness was always favored in small clusters. Together, selection both within-and between natural clusters of colonies was detectable, but highly contingent on the number of competing colonies.

---

Natural selection is context-dependent. In social organisms, context-dependent selection in individual-level traits forms the basis of diverse and powerful evolutionary forces, like indirect genetic effects, social selection, frequency-dependent selection, and their intersections (*1-5*). We contend that the context-dependent nature of selection observed in individual traits could transcend to the level of the group and their collective behavior too. For instance, the collective traits that enable group success might depend on the presence and phenotypes of surrounding groups, resulting in cases where the phenotypic compositions of some clusters of groups outperform other clusters. Whether selection occurs above the level of the group is not often evaluated in the multilevel selection literature (*6-9*), and the idea is often viewed critically on conceptual grounds. Field data documenting such selection in intact free-living systems are particularly rare. However, if selection above the level of the group is present, then it has the potential to change the evolution of individual and collective traits across a variety of social organisms and communities, especially in systems where social groups interact with each other intensely.

Here we evaluate how the number of neighboring rival groups changes two tiers of selection on collective behavior in a social spider (*Stegodyphus dumicola*). Specifically, we predict that the intensity of competition experienced by a focal group will scale positively with the number of rival groups nearby, which could alter selection both within and among clusters of colonies and potentially select against clustering (*10*). Prior work on *S. dumicola* showed that frequency-dependent selection can, at least in principle, act on among-group differences in collective behavior (*11*). In experimental clusters of colonies, the success of aggressive groups decreases as they become common within clusters. This is because aggressive colonies are more sensitive to low resource conditions, and prey are scarcer in clusters dominated by aggressive colonies. We therefore predicted that i) increasing competition between colonies (i.e., increasing the number of rivals) will reduce the performance of aggressive colonies, and ii) the performance of aggressive clusters of colonies will decrease as the number of colonies in the cluster increases.

To examine the degree of selection on collective behavior within and among clusters of spider colonies, we monitored the foraging phenotypes and performance of colonies from March 2018 to March 2019. Natural clusters of colonies (1-9 colonies, Fig. 1a) were identified along road side fences February-March 2018 in South Africa. We evaluated the collective foraging aggressiveness and colony size of each colony within each cluster. Foraging aggressiveness was evaluated thrice over two days in 2018 and 2019 by counting the number of individuals that responded to simulated prey item in the web. Starting colony sizes were estimated using the volume of the colony’s nest (i.e., a disk-shaped cylinder), which is tightly associated with the number of spiders in the colony for fence-dwelling colonies (r^2^ = 0.72-0.85, varying slightly among years). Variation in collective foraging aggressiveness is repeatable within and across generations in *S. dumicola*, and colony differences are transmitted from parent to daughter colonies during fission events, which create local clusters of related colonies upon which selection could act (*12*). This parent-offspring resemblance is itself retained across years (*11*). Thus, this system fulfills the rare pre-conditions necessary for a phenotypic response to selection both within and among clusters of colonies.

**Figure 1:**
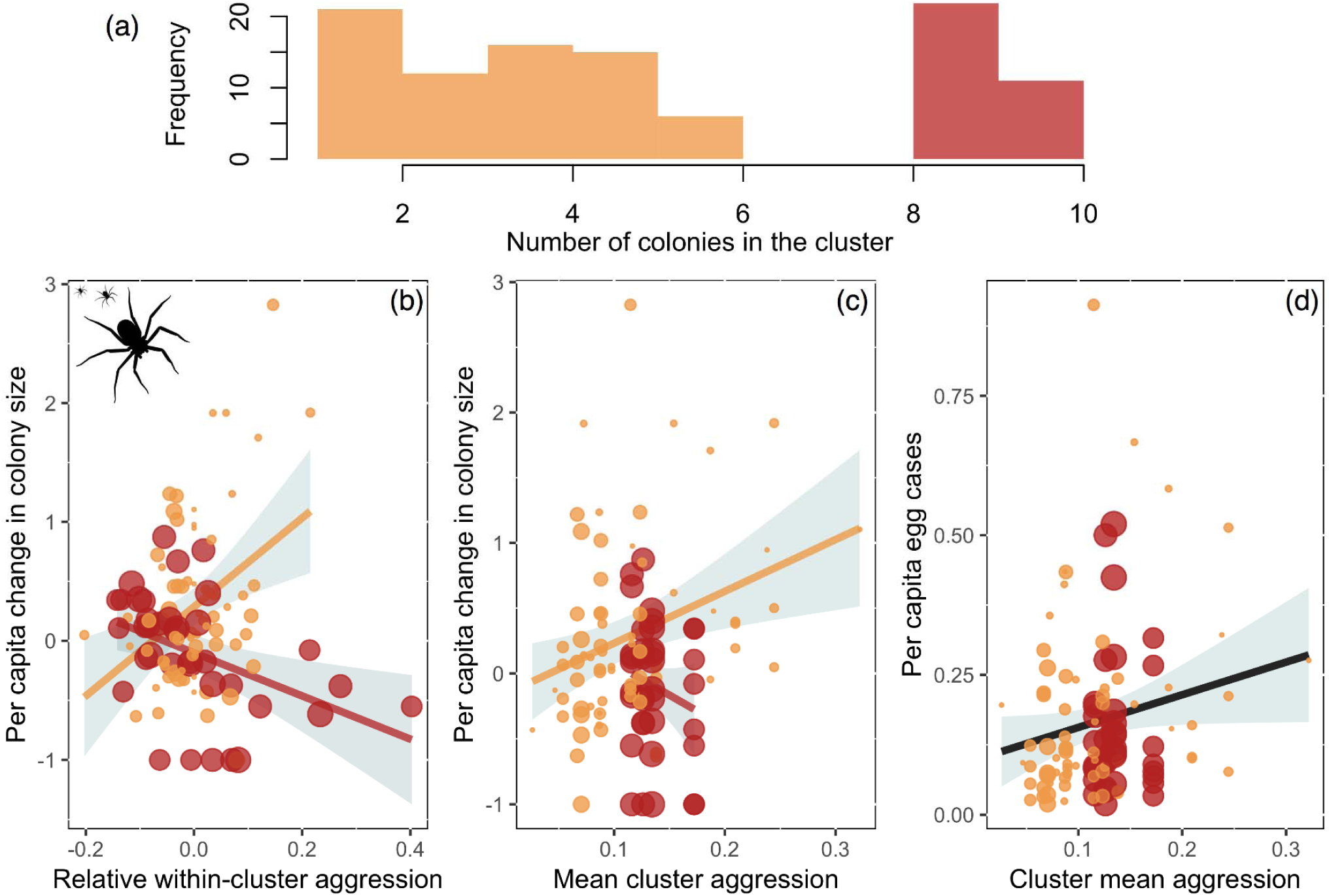

Colony success was determined at two time points. In June 2018, when colonies are dormant during the Austral winter, nests were removed from their substrate and dissected at hotels to count the number of egg cases produced by the colony in 2018 (*egg case #*). Colonies were then re-adhered to their former positions using staples and aluminum wiring to supplement supporting structure. In March 2019, we destructively recollected colonies, dissected them open again, and counted the number of females within each nest. This allowed us to record the change in colony size from one generation to the next (Δ *colony size*).

Colonies in larger clusters were less successful, and this was true for both change in colony size (*main effect* = −0.171 ± 0.05; p = 0.001) and per capita egg cases (*main effect* = −0.116 ± 0.06; p = 0.05). This finding confirms prior correlative and experimental findings that colonies of social spider compete (*11, 13, 14*), and conveys that larger clusters exacerbate competition for prey resources. Thus, selection should disfavor high levels of aggregation and promote greater dispersal, especially when clusters are composed of related colonies, which is not uncommon in *S. dumicola (12)*.

Selection on within-cluster collective aggression was dependent on cluster size. In small clusters of colonies, higher aggressiveness relative to neighboring colonies was favored, whereas non-aggressive colonies were favored in large clusters. This was true both for colonies’ proportional change in size (Fig. 1b; *interaction*: −0.336 ± 0.06, LRT = 30.05, p < 0.001, *main effect*: 0.171 ± 0.057), and *per capita* egg case production (Fig. S1; *interaction*: −0.175 ± 0.036, LRT = 25.33, p < 0.001, *main effect*: 0.314 ± 0.035). Selection on aggressiveness is thus density-dependent, switching from positive in small clusters to negative in larger clusters, likely as a result of increased resource competition in large clusters because aggressive colonies are known to outcompete docile colonies if resources are plentiful (*11*). Density-dependent social or multilevel selection has been observed in both plant and animal systems (*15-18*). However, the findings here are somewhat unique, first because the direction of selection is completely reversed in a density-dependent manner and not simply magnified, and second, because the density-dependence occurs at the level of the colony and their collective behavior rather than individual level traits.

We detected selection among clusters of colonies as well. Clusters of aggressive colonies grew proportionally more if the cluster was small, while this pattern was reversed in large clusters (Fig 1c; *interaction*: −0.161 ± 0.074, LRT = 4.655, p = 0.031, *main effect*: 0.006 ± 0.075). For *per capita* egg cases production, however, more aggressive clusters were always more successful, regardless of cluster size (Fig. 1d; *interaction*: −0.045 ± 0.095, LRT = 0.261, p = 0.61, main effect = 0.149, se = 0.056, LRT = 25.334, p < 0.001). Thus, the advantage accumulated by large and aggressive clusters until egg case production is possibly offset by costs following the emergence of the next generation. Alternatively, the selective advantage of large aggressive clusters could continue across generations but be hidden by increased long-distance dispersal from these clusters, potentially linked to local prey resources (*19*). The long-distance dispersal abilities of this species prevent us from discriminating between these interpretations. However, the possibility for conflicting selection within vs. among clusters at large cluster sizes cannot be ignored for now: non-aggressive colonies produce more egg cases than their many neighbors, but large neighborhoods of aggressive colonies are still more fecund in aggregate (Fig. S1). What becomes of this cluster-level advantage is unclear.

Despite more than a half-century of scientific debate on the efficacy of selection above the level of the individual and dozens of papers detecting its presence, multilevel selection remains one of the most instantly controversial topics in evolutionary biology. Here were use multilevel selection analysis to evaluate selection within and across two levels of selection, both of which occurs above the level of the individual: i) groups and ii) groups (clusters) of groups. The *S. dumicola* system is suited for such an analysis because between-group differences in collective behavior are transmitted with fidelity down colony lines, and because the differences in colony aggressiveness cannot be linearly traced back down to the phenotypes of individual constituents. Collective behavior is instead determined by a highly non-additive process that depends on keystone individuals (*20*), social network structure (*21*), and a colony’s social history (*22*). Yet, our data provide evidence that selection at the level of colonies and beyond are readily detectable in free-living colonies of this species, and that these levels of selection are potentially at odds for some cluster sizes. It therefore appears that evolutionary processes that emerge from feedback (positive or negative) between traits and the competitive environment are not restricted to individual levels traits, theoretical considerations, or contrived experimental settings. They are perhaps instead reasonably common and generalizable features of selection that scale from the level of genes to neighborhoods of competing societies, and conceivably beyond.

## Natural History

Social *Stegodyphus* live in multifemale colonies that cooperate in alloparental care, web maintenance, and cooperative hunting (*23, 24*). Colonies are usually founded what one or a few pre-mated females disperse to found a new colony, which grows by the breeding of brothers to sisters for multiple generations. Only a small proportion of males move between colonies, which results in high levels of relatedness within groups, and variable levels of relatedness between neighboring groups (*12, 23, 24*).

## Cluster Identification

We searched for colonies of *S. dumicola* positioned along fences (highways R74 and R714) by driving long stretches of highway. Clusters were defined as groups as nests that resided within 1m of another nest, but where the majority of the capture web was not shared with other nests. We focused on fence-dwelling colonies because 1) colonies on fences outperform tree-dwelling colonies, 2) the largest and densest clusters reside on fences, and 3) fence-dwelling nests are two-dimensional (plate-shaped), which eases their dissection and reassembly (*25*). In March 2019, we destructively recollected colonies, dissected them open again, and counted the number of females within each nest (*colony size*). Some focal clusters vanished entirely during the study from unknown causes. Only clusters that were tracked both years are considered here.

## Nest Volume & Group Size

We used nest volume to estimate the starting colony sizes of the colonies in the present study. Nest volume was estimated using a cylinder formula, with the colony’s radius and depth measured in situ using standard tape measurer or digital calipers, if the colony was small enough. Using nest volume as a proxy for groups size is not uncommon in the social spider literature, and nest volume provides only a coarse approximation of colony size in social *Stegodyphus* when they do not reside on fences (r^2^ = 0.35-0.69) (*26*). Therefore, in March 2018 we estimated the volume of 37 fence-dwelling colonies and then counted the number of adult female spiders therein. We found that nest volume explained 83.5% of the variation in colony size in these colonies, and that ending nest volume explained 72% of the variation in our focal colonies here. Thus, nest volume provides a more precise estimate of colony size in fence-dwelling *S. dumicola*, presumably because fences are more homogeneous environments for colony creation and expansion.

## Collective Foraging Aggressiveness

Colonies’ foraging aggressiveness was estimated by the number of attackers deployed during a staged encounter with prey (after *11, 27*). Trials were initiated by placing a 1 cm × 1 cm square of computer paper in the capture web, and then vibrating the piece of paper for three minutes or until the spiders made contact with the paper and seized it. The paper was vibrated using a handheld vibratory device with a thin aluminum prod extending from one end. The vibrating prod was then placed gently against the paper, causing is to flitter back and forth. We then counted the number of spiders that emerged in response to this vibratory stimulus. Colony foraging aggressiveness was evaluated thrice in 2018, twice in one day and a third time a day later. In 2019, surviving colonies were assayed 1-3 times more, to evaluate whether between-group differences in collective behavior were associated across generations, which they were (F_1,99_ = 99.52, main effect = 0.96, se ± 0.10, p < 0.0001). The majority of colonies were assayed in an identical manner to 2018. However, for a small subset of colonies, foraging aggressiveness was measured only once in 2019 because the field season was cut short owing to a medical emergency.

## Removal & Dissection

In June 2018, when colonies are dormant during the Austral winter, nests were removed from their substrate and dissected at hotels to count the number of egg cases produced by the colony in 2018 (*egg case #*). Colonies were refrigerated during this process as to keep spiders inactive prior to and after the dissection process. The perimeter of the nest was then stapled closed and colonies were re-adhered to their collection points using aluminum wiring to supplement their supporting structures and staples.

## Statistical Methods

To separate colony aggression into within cluster and among cluster components, we calculated the mean aggression of colonies within each cluster in 2018 (giving “cluster average aggression”), and then subtracted this from each colony’s mean aggression score in 2018 (giving “colony Δ aggression”). This approach for separating levels of selection is known as “contextual analysis” (*28, 29*). We then entered each of these terms into a linear mixed-effect model with a Gaussian error distribution, with the proportion increase in size of the colony between 2018 and 2019 as the response variable. For this measure of colony success, colonies that went extinct have a score of 0, while colonies that doubled in size have a score of 1, scores ranged from 0 to 2.826. The effect of colony Δ aggression corresponds to within-cluster selection, while the effect of cluster average aggression corresponds to among-cluster selection. We added the number of colonies in the cluster as a fixed effect and interacted this term with both cluster average aggression and colony delta aggression, to determine if either level of selection was density dependent. All three of these fixed effects were mean-centered and divided by their own standard deviation, meaning each variable has a mean of 0 and a variance of 1, making the regression coefficients easier to interpret (*30*). Cluster ID was entered as a random effect. We tested each interaction term for significance using likelihood ratio-tests, and if it was not significant at α = 0.05, we removed it and tested the main effect in the same manner.

To confirm our findings were robust to the choice of colony performance metric, we re-fitted this model with the number of egg-cases in the colony as a response variable, an offset of the log of colony size in 2018 (effectively modelling egg cases per capita) and a Poisson error distribution. Models were fitted in R 3.5.3 (*31*) using the package “glmmTMB” (*32*).

**Figure S1:**
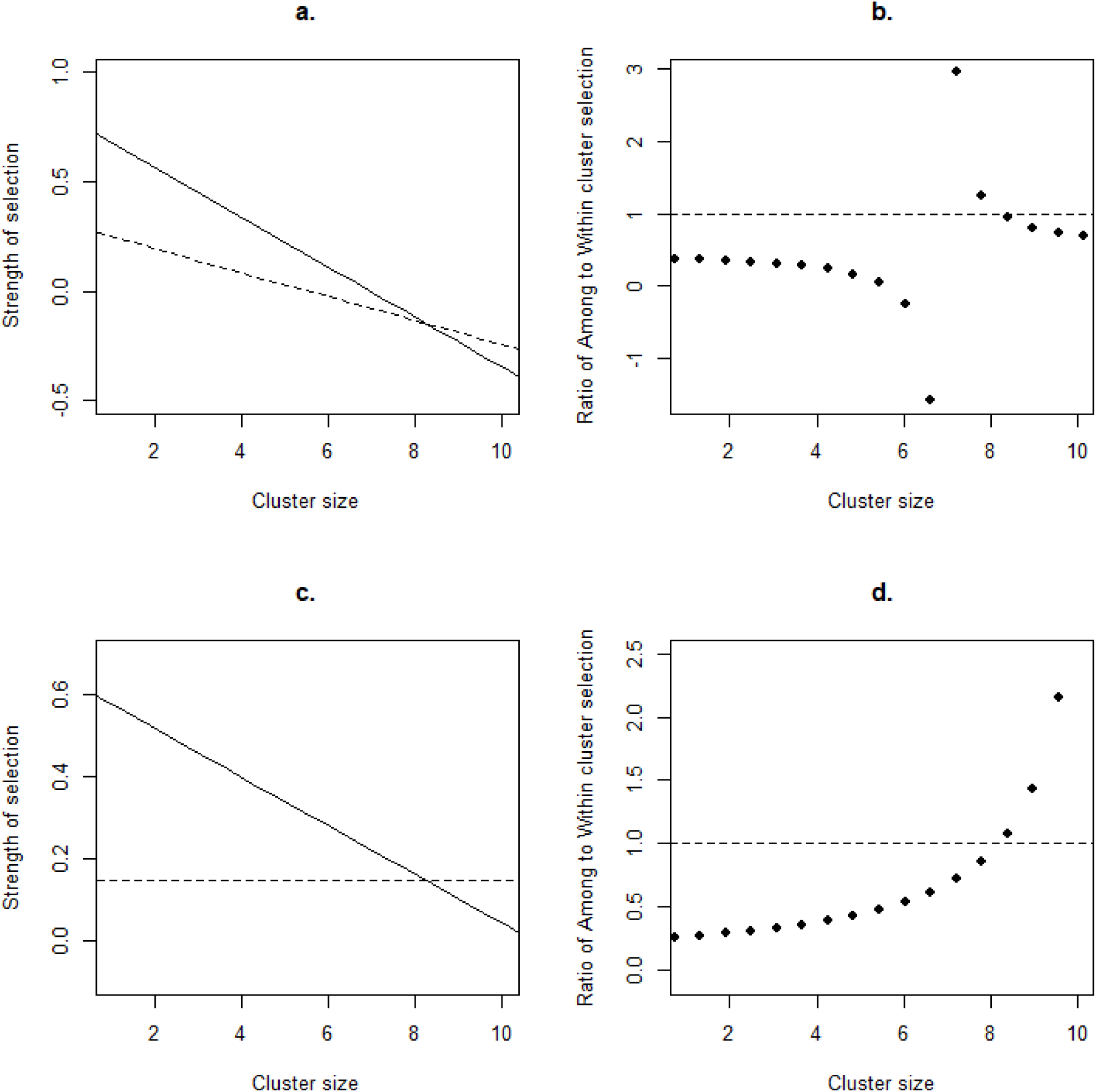
Top panels (a & b) depict selection estimates based on change in colony size, and the bottom panels (c & d) depict selection based on egg case production. The left panel (a & c) depicts the strength of within (solid line) and among (dashed line) cluster selection at various cluster sizes. The right panel (b & d) is how the ratio (among/within) changes with cluster size.

